# Psoriasin inhibits microbial growth in food by metal sequestering

**DOI:** 10.1101/2025.07.20.665715

**Authors:** Meital Gitler, Guy Sivan, Yoav Breuer, Chieh Chang, Marina Sova, Zvi Hayouka, Maayan Gal

**Author notes:** **Corresponding author** Maayan Gal.

## Abstract

Food spoilage is a significant economic and environmental concern, and it is estimated that ∼30% of fresh food is destroyed due to food spoilage between harvest to consumer. Current food preservatives are chemicals that are associated with various health risks and often have limited effectiveness under certain conditions like pH and temperature. Consequently, there’s a growing need to develop effective, natural, and economical food preservatives. Herein, we studied the natural protein psoriasin as a potential food preservative. Psoriasin is naturally secreted in the oral cavity and has an effective and validated antimicrobial activity which makes it a potentially effective and safe protein-based food preservative. Indeed, our preliminary results show promising antimicrobial activity of the recombinant protein against food-related microbial organisms in vitro and in various food types. In addition, it is recombinantly expressed at high levels, which could set a cost-effective manufacturing process. These results set psoriasin as a safe and effective natural food preservative, addressing consumer demand for healthier food options and reducing food waste.

## Introduction

### Food waste is a major environmental and economic concern

According to the Food and Agriculture Organization (FAO), about 1.3 billion tons of food are lost during the different stages of food production [1-3]. The high food loss requires intensive labor during production, leading to an increasing carbon footprint and a significant economic burden. Indeed, it is estimated that in Europe alone, food waste is worth €143 billion per year and is responsible for 15% of all greenhouse gas emissions [4-7]. Food spoilage is also a major cause of foodborne diseases, a major growing concern according to the Center for Disease Control and Prevention (CDC) [9,10], with millions of people sick due to foodborne illnesses [7-10].

### Microbiological sources leading to food spoilage

The primary cause of food spoilage is microbial contamination, mainly by bacteria and fungi [11,12]. For instance, poultry is among the main food types whose spoilage is driven by bacteria such as Salmonella and Campylobacter. Pseudomonas fluorescent and non-fluorescent strains are prevalent bacterial species leading to spoilage [13]. In red meat and seafood, bacterial species responsible for spoilage are Acinetobacter, Pseudomonas, and lactic acid bacteria [14,15]. Another major spoilage source is fungal contamination, including molds and yeasts. The main species associated with food spoilage are Fusarium, Aspergillus, Alternaria, and Penicillium strains [16-18]. Fungal contamination is associated with up to 10% of food product waste and up to 30% of crop losses [19-21]. Thus, solutions capable of preventing microbial growth that are safe for the environment and consumers are required.

### The protein psoriasin has promising antimicrobial activity as a food preservative

The food industry is turning towards “clean label” food products free from synthetic preservatives. However, only a limited number of natural and efficient materials are approved for food. Natural proteins offer a great alternative to prevent spoilage caused by microbial contamination and are used in various food-related applications. Thus, peptides and proteins are attractive natural agents for the development of antimicrobial preservatives in food. The S100 protein family is expressed in vertebrates and plays a significant role in a broad range of cellular signaling processes [22]. With more than 20 members, these proteins are characterized by a relatively low molecular weight of about 10-12 kDa, comprising two calcium-binding EF motifs. As a major Ca^2+^ signaling transporter, the S100 proteins are involved in various pathologies [23-26]. Indeed, dysregulation of S100 proteins is implicated in cancer, neurodegenerative disorders, and chronic inflammatory conditions [27-34]. Several members of the S100 family participated in immunity against microbial pathogens. These include the S100A7, also known as psoriasin, and the S100A12, also known as Calgranulin C. The S100A7 has roles in immune responses and wound healing [35]. Psoriasin possesses effective antibacterial and antifungal properties [36], and studies have shown it can disrupt fungal cell membranes, leading to fungal death [37], and prevent the adhesion of *C. albicans* on PMMA-based dentures [38]. Moreover, previous studies have demonstrated that psoriasin exhibits effective antimicrobial activity [39,40]. It also exerts potent antibacterial activity against *Escherichia coli* through high-affinity Zn^2^+ binding [41]. Additionally, S100A7 functions as a novel antifungal adhesin by binding β-glucan in the *C. albicans* cell wall, inhibiting fungal adhesion and enhancing epithelial cytokine responses [37].It was shown that S100A7 predominantly exists as a homodimer stabilized by calcium binding, with each monomer contributing two His_3_Asp motifs at the dimer interface that coordinate Zn^2^+ binding [42]. The dimeric form is also correlated with its antibacterial activity [41,43,44]. Based on its natural occurrence within the oral cavity and effective antimicrobial properties, we investigated the potential of psoriasin as a natural antimicrobial food preservative.

## Results

In the first step, we evaluated the ability of psoriasin to inhibit the growth of various food-related microbial strains. The strains were inoculated into Tryptic Soy Broth (TSB) media and incubated at 30 °C and the optical density at 600nm was monitored. **Figure 1** shows the growth curves of various strains with variable psoriasin concentrations and non-treated (red curve) as a control. Psoriasin concentration of 0.3% w/v inhibited completely inhibited *E. coli, L. Lactis, E. Cloacae* and partially inhibited *L. Sakei, L. Monocytogenes, P. Fluorescens, S. Abony and S. Cerevisiae*.

**Figure 1.**
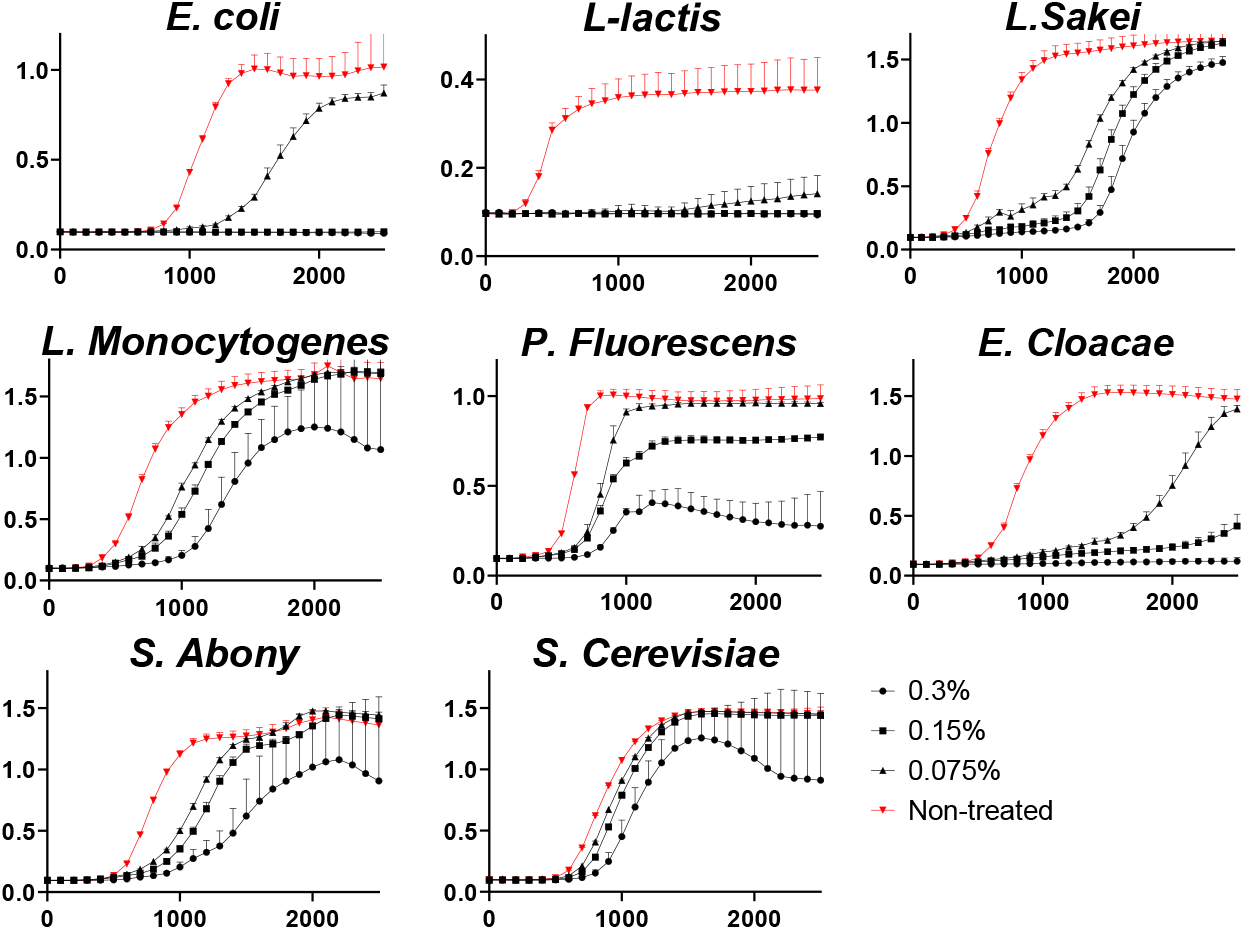
Microbial growth curves in the presence of variable psoriasin concentrations. Monitoring of microbial growth was conducted by measuring absorbance at 600 nm. Error bars represent the average of three replicates (±SD). Red curves show the growth of non-treated samples.

**Figure 2.**
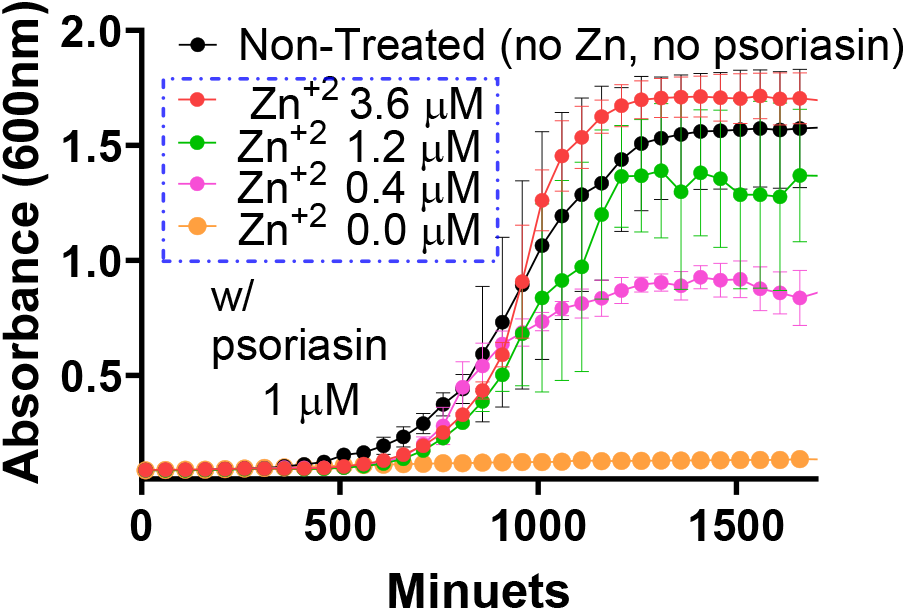
Growth curves of *C. albicans* in the presence of 1 µM psoriasin and varying zinc concentrations. Black curve – no psoriasin and no zinc were added (reference growth curve). Red, green, pink and orange – growth curves with decreasing zinc concentrations of 3.6, 1.2, 0.4 and 0 mM of ZnCl2, and in the presence of constant psoriasin concentration of 1 mM. Monitoring C. albicans growth was conducted by measuring absorbance at 600 nm. Error bars represent the average of three replicates (±SD).

We previously showed the effect of psoriasin on *C. albicans* growth in-vitro and on dentures [38]. To obtain validation for the psoriasin mechanism of action, we incubated *C. albicans* with psoriasin in the presence of variable zinc concentrations. The black curve shows the background growth in which no psoriasin or zinc was added to the medium. The orange curve shows the inhibitory effect of 1 µM psoriasin, which completely diminishes fungal growth without zinc. The pink, green and red curves show the growth of *C. albicans* in the presence of 1 µM psoriasin but with increasing zinc concentrations of 0.4, 1.2, and 3.6 µM, respectively. In the presence of psoriasin and the lack of external zinc, no growth is observed. However, the addition of zinc recovers the ability of the fungi to proliferate even in the presence of similar psoriasin concentration.

To evaluate the potential activity of psoriasin in food, we tested its effect in chickpeas (Hummus). **Figure 3** visualizes the effect psoriasin has on microbial growth in chickpeas. Hummus paste was prepared and plated in 6-well plates without and with inoculation of 500 spores of Aspergillus flavus. Samples were left at room temperature for 10 days. Whereas no microbial growth is observed in samples treated with psoriasin (top row), clear microbial growth is seen in the non-treated samples (bottom row).

**Figure 3.**
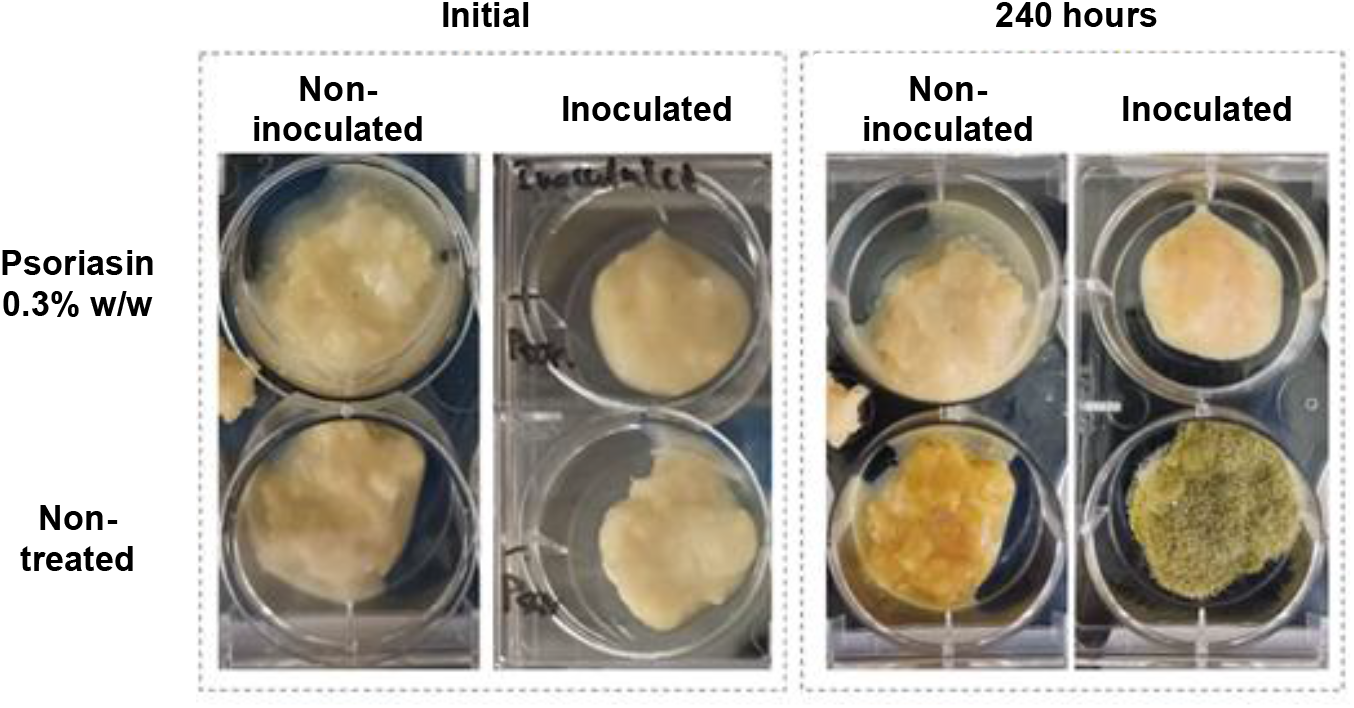
Psoriasin activity in food. Hummus without and with 0.3% w/w psoriasin at 25 C following 240 hours. Inoculated Hummus was mixed with 500 spores of Aspergillus flavus. Top row: non-treated samples. Bottom row: samples treated with 0.3% w/w psoriasin.

## Discussion

Food security is a major challenge in the current times and there is a critical need to develop preservation technologies that will lead to better utilization and reduced food waste. Of particular importance is the long-term storage and preservation of fresh food. Currently, no sufficiently effective materials are approved for use against fungi in food, and substances like potassium sorbate and LAE have limited activity and are considered undesirable for consumption.

Our study evaluates the ability of natural protein psoriasin to act as a food preservative. Unlike synthetic preservatives with potential health concerns, natural proteins offer a safer option that aligns with consumer demand for “clean-label” food products that naturally harbor broad-spectrum antimicrobial activity. Indeed, psoriasin exhibits effectiveness against both bacteria and fungi, which are the primary culprits behind food spoilage. This broad-spectrum activity offers an advantage over traditional preservatives, many of which target specific types of microbes. Moreover, the metal-chelation mechanism targets a fundamental nutritional requirement shared by bacteria and fungi, reducing the likelihood of rapid resistance development. Last, the protein is recombinantly produced with high yield even under regular lab apparatus, indicating that a potential cost-effective manufacturing is feasible using more advanced fermentation infrastructure.

Proteins could also exhibit several disadvantages. These include proteolytic and thermal instability that can inactivate it during processing or storage. In addition, potential immunogenic responses or allergenicity may occur. In the case of psoriasin, food matrices rich with divalent metals may limit its activity. Such challenges could be overcome via protein engineering approaches for the design of optimized variants with higher stability and/or zinc affinity combined with bioinformatics that could identify natural homologs harboring the required activity profile we are looking for. In addition, encapsulation technologies could protect the protein during pasteurization and provide controlled release. Future research dealing with these challenges could position natural proteins and specifically psoriasin as the next-generation protein preservatives aimed at reducing food waste and meeting consumer demand for safer, more natural products.

## Methods

### Expression and purification of recombinant psoriasin

The psoriasin gene, optimized for *E. coli* expression with an N-terminal HISx6 tag, was synthesized and cloned into the pET-28a vector (Genscript Ltd). The plasmid was transformed into *E. coli* BL21 (DE3) pLysS. Bacterial cultures grew at 37°C and 200 rpm until an OD_600_ of ∼0.8 was reached. Protein expression was induced with 1 mM IPTG for 16 hours at 25°C. Following sonication, the lysate was centrifuged at 16,000xg for 30 minutes. The supernatant was loaded onto a nickel column chromatography, and following extensive washing, the protein was subsequently eluted using 300 mM imidazole. The protein was then buffer-exchanged into phosphate-buffered saline (PBS) and frozen with 50% glycerol. To analyze the protein on SDS-PAGE, samples from each purification step were collected and resuspended in 4X Laemmli sample buffer (BioRad, Cat. #1610747) and boiled at 95°C for 10 min. Samples were subjected to 15% SDS-PAGE gel and stained with Coomassie brilliant blue (Sigma-Aldrich, Cat. #115444).

### *C. albicans* growth curves and absorbance

To monitor the effect of zinc concentration on *C. albicans* growth, 10 µl from its glycerol stock was diluted in 10 ml of RPMI-MOPS and incubated at 30°C for 12 hours. Thereafter, the optical density at 600 nm (OD_600_) was measured and adjusted to an initial optical density of 0.001 by diluting the stock into a fresh RPMI-MOPS. Subsequently, 100 µL aliquots were transferred to a transparent 96-well plate and mixed with the indicated psoriasin and zinc concentrations. The plate was incubated in a Biotek Synergy H1 microplate reader at 30°C under continuous orbital shaking and absorbance readings were taken at 600 nm from three replicate wells.To probe zinc-dependent activity, *C. albicans* was cultured as above and exposed to 1 µM psoriasin in the presence of 0, 0.4, 1.2, or 3.6 µM ZnCl_2_.

### Microbial Growth-Inhibition Assay

The following strains were purchased from Chemitec ltd. *Enterobacter cloacae, Lactococcus lactis, Saccharomyces cerevisiae, Lactobacillus sakei* and *Salmonella* abony. *Listeria monocytogenes* and *Pseudomonas fluorescens* were received from the laboratory of Livnat Afriat-Journo. *Candida albicans* and *Aspergillus flavus* were received from the laboratory of Nir Osherov at Tel Aviv University. Cultures were diluted to OD_600_ = 0.01 in fresh TSB medium and dispensed into sterile 96-well microplates containing psoriasin at indicated concentrations. Plates were incubated in a microplate reader (BioTek Epoch 2) at 30 °C with orbital shaking; OD_600_ was recorded in triplicate.

### Hummus Challenge Test

Frozen commercial hummus (100 g) was thawed and boiled for 1 hour to inactivate background microbiota. The cooked paste was homogenized and supplemented with tetracycline (10 µg mL^−1^) and chloramphenicol (25 µg mL^−1^). The treated hummus was divided into four 10 g aliquots and combined with 7 mL of the indicated additives to obtain: (i) 0.3% w/w psoriasin without fungal inoculation; (ii) 0.3% psoriasin with 500 *A. flavus* spores; (iii) uninoculated control; and (iv) inoculated control containing 500 *A. flavus* spores. Samples were stored at 25 °C for 10 days.

## Acknowledgments

The research was funded by the Israeli Ministry for Science, Technology and Space (Grant# 000735), Bountica Ltd, and the Marian Gertner Institute for Medical Nano-Systems.

